# Walking selectively modulates behaviorally relevant auditory change responses in humans

**DOI:** 10.1101/2025.08.05.668646

**Authors:** Ishrat mehmood, Haoxuan Xu, Zhouyou Dai, Cheng Chang, Peirun Song, Yuying Zhai, Xuehui Bao, Hangting Ye, Nayaab Shahir Pandit, Sirou Guo, Yanyan Wang, Lingling Zhang, Pei Chen, Wanshun Wen, Xuan Zhao, Gang Chen, Matthew J. Davidson, David Alais, Xiongjie Yu

**Author notes:** Corresponding author (X.Y.). These authors contributed equally (Ishrat mehmood, Haoxuan Xu).

## Abstract

Perception in natural environments unfolds while the body is in motion, yet how locomotion shapes cortical computations for fine-grained and dynamic scene analysis remains unclear. Here, we combined over-ground walking with mobile EEG and a transitional click-train paradigm to test how natural locomotion influences auditory processing in humans. Participants listened to two-segment click trains in which a subtle inter-click-interval (ICI) change signaled a pitch shift and performed a task-engaged change-detection task while either sitting or walking; an independent cohort also completed a passive-listening version. Walking elevated behavioral detection thresholds and slowed responses, indicating reduced sensitivity to fine temporal changes. Neurally, locomotion produced a response-component-specific attenuation: activity time-locked to the ICI transition—the change response previously linked to temporally integrative updating of an ongoing auditory object—was strongly reduced, whereas the offset response was largely preserved and onset responses were only modestly affected. Critically, this modulation persisted during passive listening and was selectively amplified near the perceptual threshold during task-engaged detection, revealing an interaction between motor state and cognitive demand. Thus, natural locomotion does not uniformly degrade hearing; instead, it preferentially down-weights a behaviorally critical, temporally demanding stage of auditory processing, offering a framework for understanding how motor–cognitive state shapes auditory scene analysis in the moving brain.

## Introduction

Perception in natural environments is inherently active: movements continuously reshape sensory input and couple motor and sensory systems into closed loops that support predicting, filtering and prioritizing behaviorally relevant information (Schroeder and Lakatos 2009, Morillon and Baillet 2017). Locomotion represents a particularly powerful state change. Each step modifies body geometry, vestibular and proprioceptive feedback, and arousal, while imposing concurrent demands on balance and navigation. In vision, locomotion reorganizes cortical dynamics and sensory gain, with visually evoked responses and detection thresholds modulated by both running speed and the phase of the stride cycle (Niell and Stryker 2010, Vinck, Batista-Brito et al. 2015, Davidson, Verstraten et al. 2024). By contrast, far less is known about how natural walking shapes auditory processing when listeners must monitor fine-grained temporal structure to guide behavior. Recent rodent and human studies show that sound-evoked activity in auditory cortex reflects both sound and movement rather than being uniformly suppressed (De Vos, Gandras et al. 2014, Yokota, Tanaka et al. 2017, Li, Wang et al. 2021, Robles, Kuziek et al. 2021, Swerdloff and Hargrove 2023, Vivaldo, Lee et al. 2023), yet most studies use simple oddball or steady-state paradigms and treat the auditory response as a unitary signal, even though extensive work has revealed rich deviance-sensitive coding throughout the auditory pathway (Xu, Yu et al. 2014, Zhai, Sun et al. 2019, Zhai, Auksztulewicz et al. 2020, Gong, Zhai et al. 2022, Song, Zhai et al. 2023). As a result, whether locomotion exerts a global, nonspecific impact on auditory encoding or selectively targets computationally demanding stages of processing remains a key unanswered question.

A central computation for real-world audition is temporal integration: the transformation of discrete acoustic events into coherent perceptual objects. When clicks are presented at inter-click intervals (ICIs) shorter than ∼30 ms, listeners no longer hear individual events but a single sound with a pitch determined by repetition rate (Yost 1996, Krumbholz, Patterson et al. 2003, Gutschalk, Patterson et al. 2004, Steinmann and Gutschalk 2012). Transitional click trains, in which two regular trains with slightly different ICIs are concatenated without a gap, exploit this property: at short ICIs, a subtle physical change yields a clear perceptual pitch shift. Recent cross-species work with such stimuli has identified partially dissociable neural components. In primate auditory cortex, a “change response” time-locked to the ICI transition closely tracks perceptual detection of small temporal shifts and has been interpreted as a signature of temporal merging (Song, Xu et al. 2024, Song, Xu et al. 2025, Xu, Huang et al. 2025). Offset responses to click trains encode an integrative summary of duration and temporal structure, suggesting that offset activity reflects accumulated temporal context (Song, Xu et al. 2024), while hierarchical recordings along the auditory pathway further indicate progressive temporal integration from midbrain to cortex (Ye, Song et al. 2025). Together, these findings delineate three stages: an onset response reflecting initial sound detection, a change response linked to temporally integrative updating of the auditory object, and an offset response encoding the integrated stimulus history. Importantly, onset and offset responses are relatively robust to task demands, whereas the change response is strongly linked to task-engaged change detection and likely recruits top-down, attentional inference (Asilador and Llano 2021, Song, Xu et al. 2024, Song, Xu et al. 2025, Xu, Huang et al. 2025, Ye, Song et al. 2025).

Here we leverage this mechanistic handle on temporal integration to test how locomotion reshapes auditory processing. Using mobile electroencephalogram (EEG), participants performed a transitional click-train task-engaged change-detection task while sitting or walking naturally, allowing us to decompose the auditory response into onset, change, and offset components and to relate each to behavioral performance. A separate passive-listening experiment dissociated the effects of locomotion per se from those of task engagement. This design addresses two key questions with high clinical relevance. First, does walking globally suppress auditory responses, or does it selectively attenuate the computationally demanding, change response linked to temporal integration? Second, are locomotion-related effects largely automatic, or are they amplified when listeners actively engage in demanding temporal discrimination near the perceptual threshold? As we show below, natural walking does not indiscriminately degrade auditory encoding; instead, it selectively attenuates behaviorally relevant change responses while leaving onset and offset responses comparatively preserved. These findings identify a specific computational stage through which locomotion interacts with auditory processing and provide a framework for understanding how the brain re-prioritizes resources to maintains auditory scene analysis in the midst of self-generated movement.

## Results

### Walking elevates thresholds for detecting subtle temporal changes

We first asked how natural walking affects the ability to detect small changes in temporal structure. Participants performed a task-engaged change-detection task on transitional click trains while either sitting or walking over-ground (Fig. 1a & Fig. 1b). Each stimulus consisted of two concatenated 1-s regular click trains: Train 1 had a fixed 4-ms ICI, whereas Train 2 had ICIs that differed from 4-ms by 0, 0.25, 0.5, 0.75, or 1.5% (e.g., 4.06-ms in the Reg_4–4.06_ condition; Fig. 1b). After the 2-s stimulus, a 100-ms, 1-kHz tone was presented as a cue, after which participants had 2-s to report whether they perceived a change via keyboard press.

**Fig. 1.**
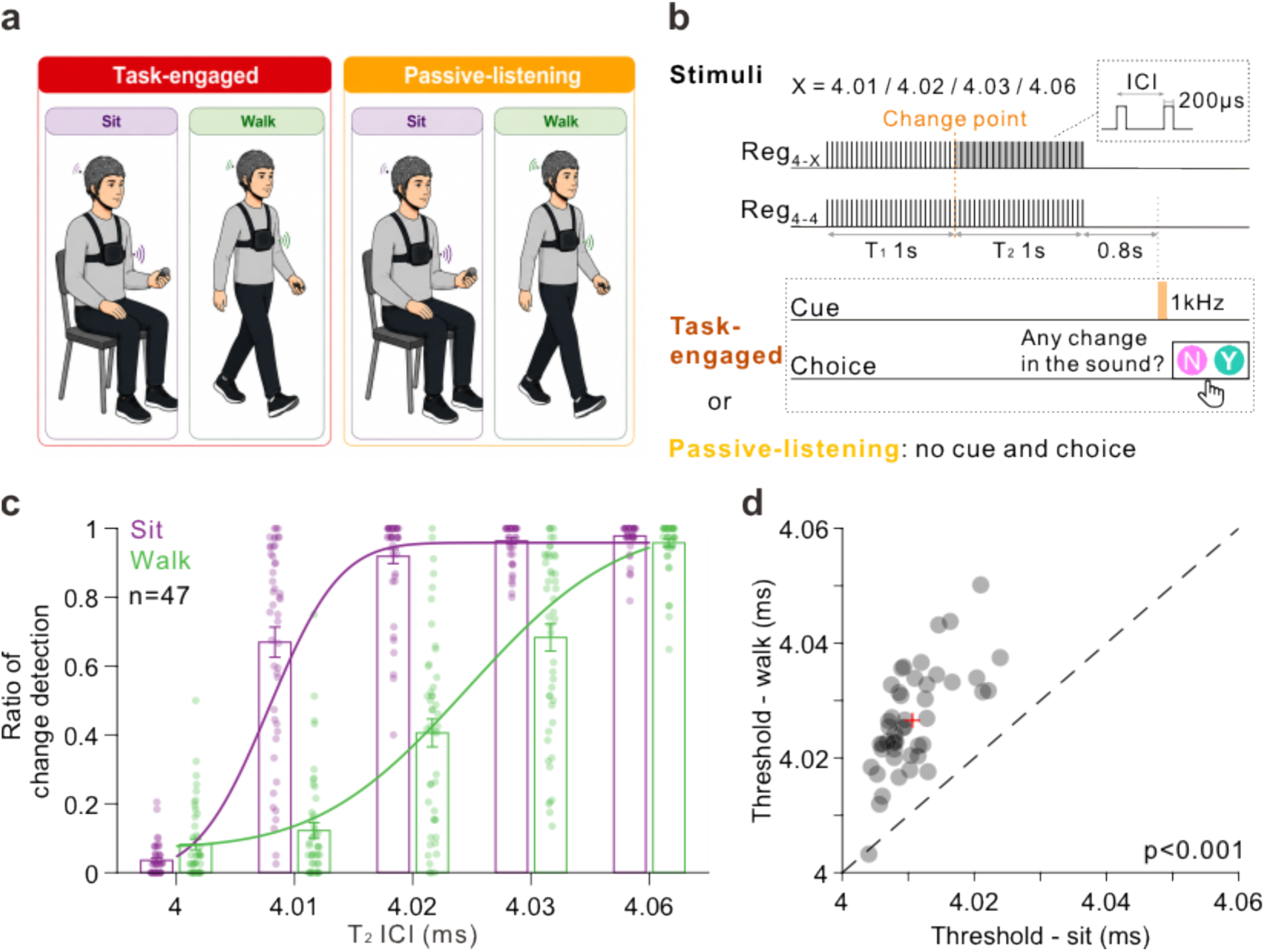
Experimental setup and behavioral change-detection performance during sitting and walking. **(a)** Schematic of the experimental setup. In the task-engaged change-detection experiment, participants performed the auditory change-detection task either while sitting (left) or walking (right), wearing a wireless EEG cap and holding a wireless response button. In the passive-listening experiment, participants completed the sessions without holding the response controller to clearly distinguish the task-free state. Neural activity was recorded during both motor states (walking and sitting) across experiments. **(b)** Structure of the transitional click-train stimuli. Two click trains composed of 0.2-ms monopolar pulses with different ICIs were seamlessly concatenated to form a transitional click train. For example, a transition from 4 ms (train 1, T1) to 4.06 ms (train 2, T2) is denoted as Reg_4–4.06_. Five stimulus types (Reg_4–4_, Reg_4–4.01_, Reg_4–4.02_, Reg_4–4.03_, and Reg_4–4.06_) were presented in random order. An auditory cue (1 kHz, 100 ms) was delivered 800 ms after the end of the click train, prompting participants to report whether a change had occurred via button press within a 2-s response window. In the passive-listening experiment, no behavioral response was required. **(c)** Change-detection rate as a function of T2 ICIs during sitting (purple) and walking (green) conditions (mean ± S.E.M.; *n* = 47). Transparent dots represent individual participants. **(d)** Scatter plot comparing change-detection thresholds between sitting and walking conditions (*n* = 47). Each dot represents one participant; the red cross indicates the group mean. Statistical significance was assessed using a two-tailed paired *t*-test (*p* < 0.001).

Psychometric functions showed a robust locomotion effect (Fig. 1c). Across participants (*n* = 47), detection thresholds were systematically higher during walking than sitting (paired *t*-test, *t*(46) = 16.36, *p* < 0.001; Fig. 1d), corresponding to rightward-shifted psychometric curves in the walking condition. Thus, even though the physical stimuli were identical, natural walking elevated the threshold for detecting subtle temporal changes in the click trains.

### Walking selectively attenuates cortical change responses

We next examined whether the behavioral impairment during walking was mirrored in cortical responses. Transitional click trains reliably evoked three temporally distinct components in the EEG: an onset response following sound onset, a change response time-locked to the ICI transition, and an offset response following stimulus termination (Fig. 2a).

**Fig. 2.**
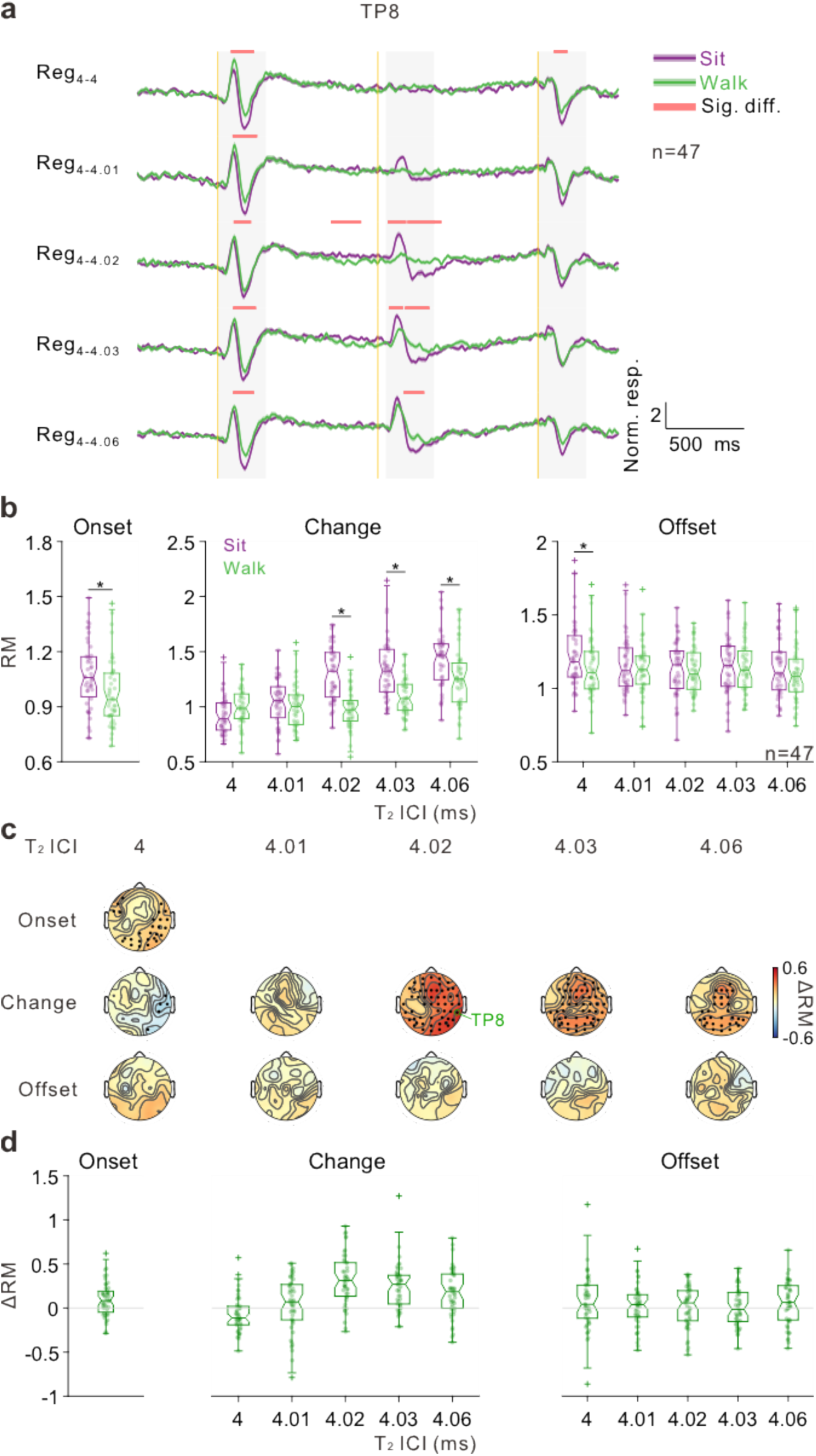
Walking selectively attenuates auditory onset and change responses. **(a)** Grand-averaged normalized ERPs at the representative electrode TP8 during sitting (purple) and walking (green) conditions (*n* = 47). Shaded gray regions indicate the predefined analysis windows for onset, change, and offset responses. Red bars at the top denote time periods showing significant differences between sitting and walking (*p* < 0.05, permutation test). Vertical orange lines mark the onset (0 ms), change point (1000 ms), and offset (2000 ms) of the transitional click train. **(b)** Response magnitude (RM; root mean square amplitude) for onset, change, and offset responses across the five T_2_ ICI conditions (Reg_4–4_ to Reg_4–4.06_), shown separately for sitting (purple) and walking (green). Each dot represents one participant (crosses indicate outliers). Notched boxplots display the median (center line), interquartile range (box: 25^th^–75^th^ percentiles), and data spread (whiskers extend to 1.5 × the interquartile range); notches indicate the 95% confidence interval of the median. Asterisks denote significant differences between sitting and walking conditions (*p* < 0.05, two-tailed paired *t*-test). **(c)** Scalp topographies of the difference in response magnitude (ΔRM = sit − walk) for onset, change, and offset responses across T_2_ ICI conditions. Dots indicate electrodes showing significant differences (*p* < 0.05, FDR-corrected, two-tailed paired *t*-test). The representative electrode TP8 is marked with a green circle. **(d)** Distribution of ΔRM (sit − walk) for onset, change, and offset responses across T_2_ ICI conditions. Each dot represents one participant.

Previous work with similar stimuli has linked the behaviorally relevant change response to dynamic temporal integration and object updating and the offset response to the integrated history of the stimulus (Song, Xu et al. 2024, Xu, Huang et al. 2025).

At a representative parietal electrode (TP8), selected because it exhibited the largest change-response amplitudes across all channels, onset responses were broadly similar across conditions but showed modestly larger amplitudes during sitting in the 95–217 ms post-onset window (paired *t*-test, permutation corrected, *p* < 0.05). In contrast, change responses were markedly attenuated during walking, particularly 67–385 ms after the ICI transition (Reg_4–4.02_, paired *t*-test, permutation corrected, *p* < 0.05), whereas offset responses showed no reliable difference between sitting and walking. Thus, locomotion does not produce a global, uniform suppression across all components; rather, it preferentially weakens the integrative, change-related activity while largely sparing termination-related responses.

We quantified these effects using a response magnitude (RM) measure for each component and condition (Fig. 2b). For the onset response, pooling across all stimuli (Train 1 always Reg_4_), RM was significantly reduced in walking compared to sitting (*t*(46) = 3.52, *p* < 0.001, Cohen’s *d* = 0.51). For the change response, locomotion induced pronounced attenuation at intermediate and high contrasts, Reg_4–4.02_, Reg_4–4.03_, and Reg_4–4.06_ (all *p* < 0.001), whereas the offset response showed no significant difference between sitting and walking at any contrast.

Topographical maps of sit – walk RM differences revealed distinct spatial signatures (Fig. 2c). Onset modulation involved 27 FDR-corrected electrodes (*p* < 0.05), concentrated over bilateral parietal sites. Change-response modulation was more extensive and strongly contrast-dependent: minimal at low contrasts (Reg_4–4_, Reg_4–4.01_), peaking at Reg_4–4.02_ (54 significant electrodes), and remaining widespread at Reg_4–4.03_ and Reg_4–4.06_ (32 significant electrodes at the highest contrast). In contrast, offset responses exhibited negligible modulation, with no electrodes surviving correction across contrasts. Thus, although both onset and change responses are attenuated by walking, the effect is strongest and most spatially extensive for the change response, whereas the offset component is largely preserved.

To examine how locomotion-related modulation depends on stimulus contrast, we computed ΔRM (sit – walk) averaged across all channels as a function of ICI contrast (Fig. 2d). This all-channel average was employed to provide a robust, low-noise metric of the overall modulation magnitude, consistent with the widespread significant clusters identified topographically. For the change response, ΔRM increased from Reg_4–4_ to Reg_4–4.02_ and then slightly decreased toward Reg_4–4.06_, yielding a non-monotonic profile (one-way ANOVA, *F*(4,230) = 16.09, *p* < 0.001, *η*^2^ = 0.22). Critically, at Reg_4–4.02_the contrast near perceptual threshold ΔRM for the change response exceeded that for the onset response, indicating that locomotion exerts its largest impact precisely where change detection is most demanding. In contrast, ΔRM for the offset response did not vary across contrasts (*F*(4,230) = 0.66, *p* = 0.62, *η*^2^ = 0.01; *BF*_10_ = 0.01). Furthermore, the Bayes Factor (*BF*_10_ = 0.01provided strong evidence for the null hypothesis (i.e., no difference in ΔRM across contrasts for the offset response), rigorously confirming the selective nature of the effect. Overall, the data reveal a graded, response-type-specific pattern: modest attenuation for onset, pronounced and behaviorally aligned attenuation for change responses, and minimal impact on offset. Importantly, the walking-related effect was component-specific rather than global: offset responses recorded in the same trials and preprocessing pipeline were largely preserved, arguing against a simple explanation based on broadband movement artifacts or generalized signal loss during walking.

### Locomotion-induced modulation persists without task engagement

The above experiments were performed under a task-engaged change detection task, raising the question of whether locomotion-related modulation of the change response depends on task engagement. We therefore conducted a passive listening control experiment in an independent cohort, in which participants alternated between sitting and walking while hearing the same transitional click trains but performing no behavioral task (Fig. 1a & Fig. 1b).

At the population level (*n* = 21), Onset RM was significantly reduced during walking (Wilcoxon signed-rank test, *Z* = 2.69, *p* < 0.01, *r* = 0.59). For the change response, locomotion-induced attenuation emerged at higher contrasts (Reg_4–4.02_, Reg_4–4.03_, Reg_4–4.06_; all *p* < 0.01). The offset response showed only a small, isolated effect at Reg_4–4.01_ (*Z* = 2.83, *p* < 0.01, *r* = 0.62) with no systematic pattern across contrasts (Fig. 3a).

**Fig. 3.**
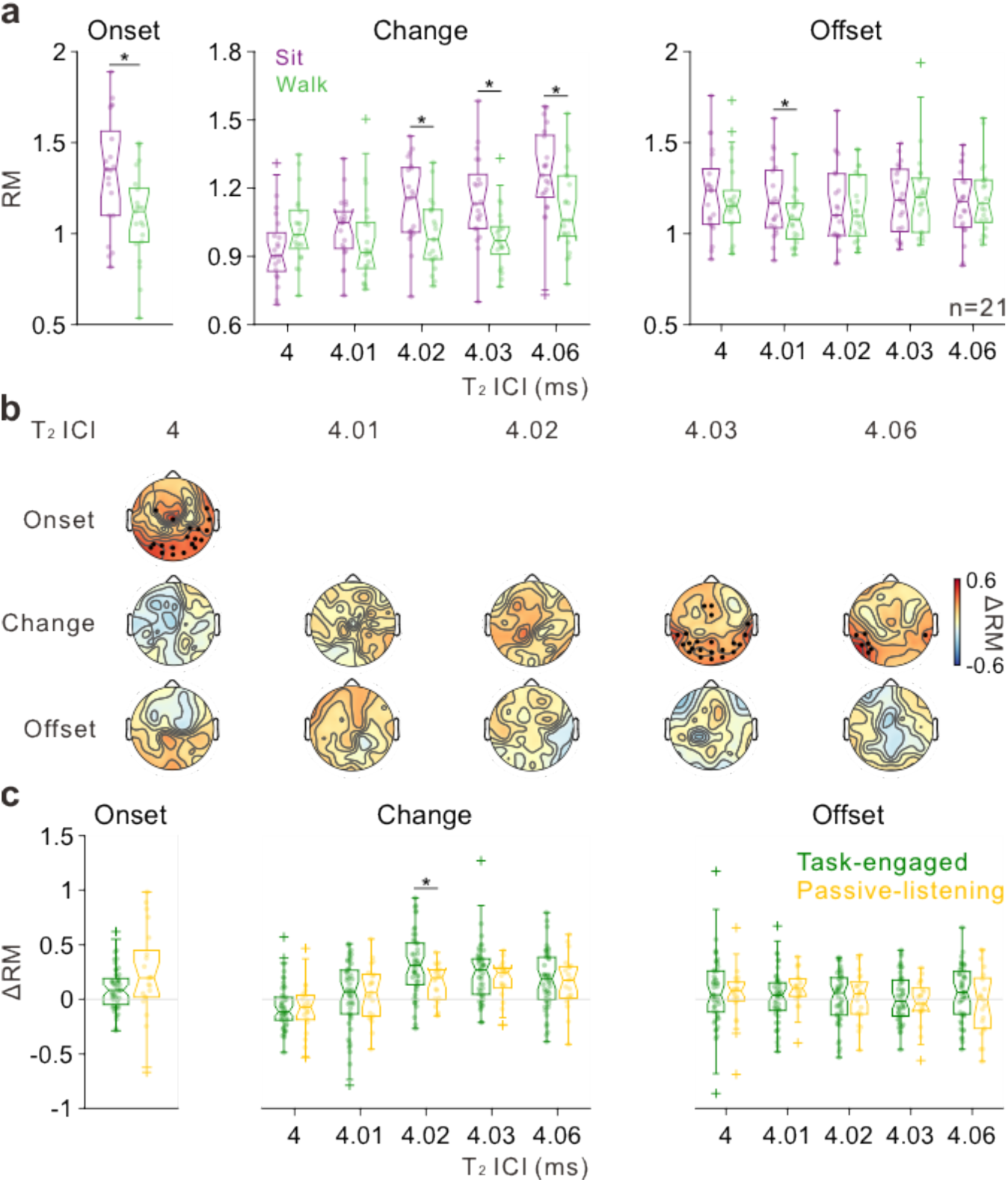
Walking-induced modulation of auditory responses during passive-listening. **(a)** Response magnitude (RM; root mean square amplitude) for onset, change, and offset responses across the five T_2_ ICI conditions during passive-listening, shown separately for sitting (purple) and walking (green). Plotting conventions are identical to those in Figure 2b. Asterisks denote significant differences (*p* < 0.05, two-tailed paired *t*-test). Sample size: *n* = 21. **(b)** Scalp topographies of the difference in response magnitude (ΔRM = sit − walk) for onset, change, and offset responses across T_2_ ICI conditions during passive-listening. Dots indicate electrodes showing significant differences (*p* < 0.05, FDR-corrected, two-tailed paired *t*-test). **(c)** Distribution of ΔRM (sit − walk) for onset, change, and offset responses across T_2_ ICI conditions, comparing task-engaged (green) and passive-listening (yellow) conditions. Each dot represents one participant. Differences between task-engaged and passive-listening conditions were assessed using a two-tailed Mann–Whitney *U* test (*p* < 0.05), reflecting the use of independent participant samples across experiments.

Topographical maps of ΔRM (sit - walk; Fig. 3b) showed fewer significant electrodes than during the task-engaged change detection task: 21 electrodes for onset over parietal–occipital sites, and contrast-dependent clusters for the change response (none at low contrast, 23 at Reg_4–4.03_, 5 at Reg_4–4.06_), with no significant offset clusters. These findings indicate that locomotion-driven modulation of the change response persists during passive listening but is weaker and requires larger physical contrasts to emerge. To directly test the role of task engagement, we compared ΔRM (sit – walk) between task-engaged and passive-listening conditions. For onset and offset responses, the magnitude of locomotion-induced modulation did not differ significantly between tasks, suggesting that these components are modulated by walking largely independent of behavioral demands. By contrast, ΔRM for the change response at Reg_4–4.02_ was significantly and selectively larger during the change-detection task than in non-task listening (Mann–Whitney *U* test, *Z* = 2.50, *p* < 0.05, *r* = 0.30; Fig. 3c). Thus, change responses are both locomotion-sensitive and task-augmented: walking suppresses them even in the absence of a task, and active, task-engaged change detection near perceptual threshold powerfully amplifies this modulation.

## Discussion

Perception in everyday life rarely occurs at rest: we listen and decide while our bodies are in motion. Here, we combined a transitional click–train paradigm with mobile EEG during over-ground walking to investigate how natural locomotion reshapes auditory temporal integration at distinct computational stages. We found that walking elevates thresholds for detecting subtle temporal changes (Fig. 1c–d), selectively attenuates the cortical change response far more strongly than onset or offset responses (Fig. 2a–d), and that this attenuation persists during passive listening while being synergistically amplified by active change detection, especially near threshold (Fig. 3). Together, these results identify the temporally integrative change response as a primary cortical locus where locomotion, attention, and auditory processing converge.

### Preferential Modulation of the Behavior Relevant Response

Our first main result is that walking does not uniformly down-scale auditory responses. Behaviorally, psychometric functions shift rightward during walking relative to sitting (Fig. 1c, d), indicating that listeners require larger ICI changes to detect the transition. Neurally, although both onset and change responses are reduced in amplitude during walking, but the suppression is strongest and most spatially extensive for the change response, particularly at intermediate, near-threshold contrasts, whereas the offset response is remarkably stable across motor states (Fig. 2a–d). This pattern aligns with prior work using transitional and offset-sensitive click trains: onset responses primarily reflect bottom-up detection of sound onset(Nabé, Schwartz et al. 2021), offset responses encode the termination and integrated duration-based structure of an auditory object and are relatively robust to state changes (Song, Xu et al. 2024), and change responses track temporally integrative updating of the auditory object when click-train structure changes (Song, Xu et al. 2024, Song, Xu et al. 2025).

Detecting a subtle ICI change in our paradigm requires maintaining an internal representation of the ongoing temporal pattern, integrating information across clicks, and comparing this prediction with new input operations that resemble top-down inference over temporally extended evidence (Tallal, Miller et al. 1993, Asilador and Llano 2021, Di Stefano and Spence 2025). The observation that locomotion most strongly suppresses the change response, and does so maximally at contrasts that are behaviorally near threshold (Fig. 2b, d), suggests that locomotion preferentially targets these higher-order, resource-intensive temporal computations rather than early sensory encoding. This is consistent with prior work showing that movement-related signals reconfigure network state in sensory cortices and bias sensory processing toward or away from demanding computations (Zagha, Casale et al. 2013, Parker, Brown et al. 2020, Gong, Song et al. 2024, Du, Xu et al. 2025).

Over-ground walking, although it appears automatic yet engages substantial cortical resources for balance, navigation, and multisensory integration (Stuart, Vitorio et al. 2018, Delval, Bayot et al. 2020, Rubega, Di Marco et al. 2021).Theoretical frameworks such as threaded cognition and dual-task interference propose that partially shared resources must be time-shared across concurrent operations, leading to performance decrements when demands overlap (Salvucci and Taatgen 2008, McPhee, Cheung et al. 2022, Riedel, Herzog et al. 2024). Within this framework, our data suggest that resource-intensive temporal integration and change detection are uniquely susceptible to locomotor interference than the more automatic processes supporting onset and offset responses particularly when discrimination is difficult (Fig. 2b, d). Related work in development and multisensory processing likewise indicates that temporal integration is especially vulnerable to load and state changes (Tallal, Miller et al. 1993, Dionne-Dostie, Paquette et al. 2015, Senkowski and Engel 2024). The comparison between task-engaged and passive listening directly supports this interpretation. During passive listening, the change response is still attenuated during walking (Fig. 3a–c), indicating that movement alone is sufficient to modulate temporal integration. However, locomotor attenuation is weaker and emerges only at larger physical contrasts. Crucially, when participants actively detect changes, the same physical stimuli elicit significantly stronger locomotion-induced suppression, with the largest difference at near-threshold contrasts (Fig. 3b, d), whereas onset and offset modulation are similar across tasks. This double dissociation points to a synergistic interaction between locomotion and attentional/decision-related processes recruited in demanding temporal discrimination (Proksch, Comstock et al. 2020, Liebherr, Corcoran et al. 2021, Auksztulewicz, Ödül et al. 2025). In other words, walking provides a background state shift; and active temporal comparison adds an additional layer of cognitive load. Together they selectively down-weight the most demanding auditory computation the change response while leaving offset encoding of integrated duration/structure relatively robust (Song, Xu et al. 2024).

Importantly, the behavioral impairment observed during walking is not inconsistent with a state-dependent resource-allocation account, provided that adaptation is considered at the level of whole-body behavior rather than performance in the auditory task alone. Over-ground walking imposes demands for postural control, self-motion monitoring, navigation, and the integration of visual, vestibular, and proprioceptive feedback (Stuart, Vitorio et al. 2018, Delval, Bayot et al. 2020, Parker, Brown et al. 2020, Rubega, Formaggio et al. 2021, Senkowski and Engel 2024). Under these conditions, fine-grained auditory temporal discrimination may be relatively deprioritized, consistent with resource-sharing and dual-task interference frameworks in which concurrent motor and cognitive demands compete for partially overlapping processing resources (Salvucci and Taatgen 2008, McPhee, Cheung et al. 2022, Riedel, Herzog et al. 2024). Thus, locomotion-related attenuation of the change response should not be interpreted as optimization of auditory change detection itself; rather, it may reflect state-dependent prioritization of locomotor control and broader sensorimotor demands, producing a local cost for detecting subtle ICI changes. The component-specific nature of the effect—strongest for the change response, modest for onset, and minimal for offset—is less consistent with a purely nonspecific sensory-interference account and instead supports a model in which locomotion selectively gates resource-intensive auditory computations.

Mechanistically, such preferential modulation could arise from several, not mutually exclusive, mechanisms: (i) locomotor efference copies or corollary discharge targeting the cortical layers and circuits engaged in temporal integration (Hentschke, Haiss et al. 2006, Zagha, Casale et al. 2013), (ii) neuromodulatory shifts during locomotion that preferentially affect longer-timescale recurrent dynamics (Parker, Brown et al. 2020, Senkowski and Engel 2024), or (iii) competition between internal models used for gait control and those used to predict fine-grained temporal structure in sound (Morillon, Schroeder et al. 2014, Morillon and Baillet 2017). Disentangling these possibilities/mechanisms will require invasive recordings during locomotion in animals and, where possible, intracranial recordings in humans, but the current data already constrain models by showing that dynamic updating at the moment of change not offset integration is the most vulnerable stage.

The direction of locomotion-related modulation should also be interpreted in relation to the broader active-sensing literature. Locomotion can enhance sensory gain in some systems, most prominently in visual cortex, where running increases visually evoked responses and improves visual sensitivity (Niell and Stryker 2010). However, motor-state effects are not uniformly facilitatory across modalities or cortical circuits. In the auditory system, active behavioral states and locomotion have often been associated with reduced sound-evoked responses, particularly in superficial cortical circuits, even when sensory tuning or information content is preserved (Zhou, Liang et al. 2014, Schneider and Mooney 2018). Moreover, recent work showed that auditory cortical ensembles jointly encode sound and locomotion speed during movement, while also reporting net suppression of baseline-subtracted sound-evoked responses and neural stimulus detection during locomotion (Vivaldo, Lee et al. 2023). Thus, the present attenuation of auditory responses during walking does not contradict active-sensing models; instead, it supports a more flexible view in which locomotion reweights sensory computations according to modality, behavioral context and task demands.

Our task-engaged and passive-listening experiments further clarify this interpretation. The change response was attenuated during passive listening, indicating that walking alone is sufficient to alter auditory change-related activity (Fig. 3a, b). However, this attenuation was amplified during task-engaged near-threshold discrimination (Fig. 3c), indicating an additional contribution from attentional and decision-related demands. We therefore do not interpret the effect as purely sensory interference or as purely strategic resource allocation. Rather, the data support a state-dependent gating account in which locomotion interacts with task demand to selectively modulate resource-intensive auditory computations.

### Implications and future directions

These findings have several implications for understanding hearing in natural environments. First, they argue against treating “walking” as a simple gain decrease on auditory processing: locomotion and its associated task demands selectively down-weight temporally integrative, behaviorally relevant computations while leaving onset and offset encoding relatively robust. Standard mobile EEG and brain-computer interface (BCI) paradigms that treat the auditory response as a unitary signal may therefore miss important, component-specific modulations. Second, by identifying the change response as the primary locus of locomotor modulation, our results provide a concrete neural readout for testing mechanistic models of motor–sensory interaction and temporal prediction in midbrain and cortical circuits (Morillon and Baillet 2017, Gong, Song et al. 2024, Du, Xu et al. 2025). At the same time, our approach is necessarily constrained to the gross-motor state (sitting vs walking), performed at a comfortable, self-selected speed in a controlled indoor environment. Although an additional analysis revealed no significant male–female difference in locomotion-induced modulation of the change response across ICI conditions in either the task-engaged or passive-listening experiment, the sample was not explicitly male-female balanced. Future studies with larger and more balanced cohorts will be needed to more definitively assess potential male-female differences. However, to fully characterise the coupling between movement and temporal integration, future work should systematically examine modulations by the moment-to-moment phase of the stride cycle, as well as how varying walking speed, path complexity (e.g., turning, uneven surfaces), and multitasking demands influence the generality and boundary conditions of these motor–auditory couplings. Extending the paradigm to ecologically relevant stimuli such as speech, music, or noisy backgrounds, will be essential for linking these findings to everyday listening and the broader literature on auditory–motor entrainment and beat perception (Proksch, Comstock et al. 2020, Pranjić, Braun Janzen et al. 2024, Noboa, Kertész et al. 2025). Finally, mobile EEG also lacks the spatial resolution to identify specific layers and cell types; combining similar tasks with laminar or intracranial recordings in animals during locomotion will be crucial for uncovering the microcircuits that generate the change response and its modulation.

Despite these caveats, the present data support a clear conclusion. Locomotion does not uniformly add noise to the auditory system or uniformly suppress sensory responses. Instead, it selectively reweights resource-intensive, temporally integrative and behaviorally relevant computations. This state-dependent coupling between movement and temporal integration provides a principled framework for understanding how the brain maintains auditory scene analysis while the body moves, and highlights that exploiting rather than ignoring motor state may be critical for next-generation models of active listening, mobile neurotechnology, and auditory prosthetics.

## Materials and Methods

### Experimental Procedure and Participants

This study encompassed two main experiments (Experiment 1: behavioral (Task-engaged change-detection) and neural; Experiment 2: Passive Listening), conducted in compliance with the Declaration of Helsinki (2013) (Duncan 2013) and approved by the Institutional Review Board (IRB-20230131-R). Written informed consent was obtained from all participants. All experiments were performed in a soundproof room measuring 8 m × 8 m. During walking blocks, participants walked overground back and forth along a straight 8-m path at a comfortable, self-selected pace, turning naturally at the end of each path and then continuing in the opposite direction. Walking was performed overground (i.e., not on a treadmill), and no obstacles were present. Experiment 1 (behavioral (Task-engaged change-detection task) and neural) involved 47 healthy adults (28 males, 19 females; mean age 22.98 ± 2.72 years). Experiment 2 (passive listening) included an independent cohort of 21 healthy adults (12 males, 9 females; mean age 23.90 ± 3.88 years). Thus, Experiment 1 and Experiment 2 used independent participant cohorts; sitting and walking blocks were alternated within each experiment. Participants were instructed to keep their eyes open, remain awake, listen to the sounds, and minimize unnecessary head and body movements in both sitting and walking conditions. During walking, participants were asked to maintain a comfortable walking pace and a forward-facing head/gaze posture as much as possible; strict head fixation was not required because it is incompatible with natural overground walking. In the task-engaged condition, responses were made using a wireless button press.

**Experiment 1** employed a task-engaged change detection task with alternating sitting and walking blocks to probe perceptual and neural correlates of auditory processing under different motor states. Participants completed ten alternating blocks (five sitting and five walking), with the session beginning in the sitting condition and each block lasting approximately 3 minutes. The sitting and walking blocks were alternated to temporally interleave motor-state conditions and reduce potential confounds related to fatigue, vigilance decline, habituation, learning, slow EEG drift, and time-on-task. Alternating blocks also prevented one motor state from being sampled mainly during the early or late part of the recording session, thereby helping to maintain comparable alertness and engagement across conditions. During sitting blocks, participants sat holding a wireless keypad, with auditory stimuli delivered via a chest-mounted speaker. During walking blocks, participants walked at a normal pace (∼1 m/s, ∼1 Hz stride frequency), monitored using inertial sensors, within the soundproof room with the same setup (Fig. 1a). In each trial, one of five transitional click trains (Reg_4–4_, Reg_4–4.01_, Reg_4–4.02_, Reg_4–4.03_, Reg_4–4.06_) were presented, with conditions randomized across trials (∼40 trials per condition). Each transitional click train was followed by an 800-ms delay and a 100-ms 1-kHz response cue (Fig. 1b). Participants were instructed to report whether a change occurred by pressing keys (right for change, left for no change) within 2-s following the cue. Behavioral (Fig. 1c–d) and neural (Fig. 2a–d) analyses included all click train conditions and focused on assessing sensitivity to ICI differences across motor states.

**Experiment 2** examined passive listening during sitting and walking to evaluate baseline neural responses in the absence of task engagement. Participants were presented with the same transitional click trains as in Experiment 1 (∼40 trials per condition), while no behavioral response was required. Participants were instructed to remain awake, keep their eyes open, and listen to the sounds throughout the passive-listening blocks. The experimenter monitored participants during recording and provided short breaks during recording and, when necessary, between blocks. Passive-listening blocks were also alternated across sitting and walking conditions, with each block lasting approximately 3 minutes. This design was used to avoid prolonged task-free periods and to help maintain comparable alertness, fatigue, and time-on-task across motor states. The experimental setup and stimulus delivery were otherwise identical to those used in Experiment 1. Neural analyses (Fig. 3a–c) focused on assessing sensitivity to ICI variations across motor states and comparing locomotion-related modulation between task-engaged and passive-listening conditions.

### Auditory Stimuli

All experiments were conducted in a soundproof room. Each click consisted of a 0.2-ms monopolar pulse (Fig. 1b). Regular click trains featured fixed ICIs without randomization. Transitional click trains were formed by concatenating two regular trains with differing fixed ICIs, for instance, Reg_4–4.06_ combined a 4-ms ICI train followed seamlessly by a 4.06-ms ICI train. Contrasts were defined as 0%, 0.25%, 0.5%, 0.75%, and 1.5% relative to the 4-ms baseline ICI. Stimuli were delivered through a Golden Field M23 speaker, powered by a Creative AE-7 Sound Blaster at a sampling rate of 384 kHz, and controlled via Psychtoolbox 3 in MATLAB. Sound intensity was calibrated to a constant 60 dB SPL using a ¼-inch Brüel & Kjær 4954 condenser microphone and PHOTON/RT analyzer.

### Data Acquisition

EEG data for both experiments were acquired using a 64-channel NeuSenW system (Neuracle, China), with 59 electrodes active in practice. Ground electrodes were positioned between Fpz and Fz in the frontal region. The reference electrode was placed between Cz and Pz, replacing CPz in the wireless EEG cap. Concurrent with high-density EEG signals, embedded inertial sensors captured movement data at 100 Hz, facilitating synchronized monitoring of brain activity and dynamics. EEG was sampled at 1 kHz, with electrode placement adhering to the international 10/20 system.

### Data Analysis

#### Preprocessing

Data analyses were performed using MATLAB R2021b (The MathWorks, Inc., Natick, MA, USA) and the FieldTrip toolbox (Oostenveld, Fries et al. 2011). Monopolar referencing was applied using the default reference electrode positioned between Cz and Pz. Multichannel EEG data underwent sequential preprocessing: initial bandpass filtering (two-pass Butterworth, 0.5–40 Hz), followed by epoch extraction from −1 to 3 s relative to trial onset. Independent component analysis (ICA) removed electrooculogram (EOG) artifacts. Baseline correction subtracted the mean activity from −200 to 0 ms pre-stimulus onset per trial. Motion artifacts were evaluated via a relative threshold, flagging samples outside mean ± 3 × SD as “bad.” Trials exceeding 20% bad samples were discarded as “bad trials,” and channels with >10% bad trials were labeled “bad channels.” Bad channels were excluded first (based on all-channel data), followed by bad trials from remaining channels. Event-related potentials (ERPs) were computed by averaging epochs per experimental condition, channel, and subject. Prior to group-level analysis, ERPs were normalized by dividing each channel’s data by its standard deviation per subject to minimize inter-channel variability.

During walking blocks, participants walked back and forth along an 8-m path at a comfortable, self-selected pace and turned naturally at the end of each path before continuing in the opposite direction. Because the aim of the study was to characterize auditory processing during natural overground walking rather than during isolated gait phases, brief turns at the end of the walking path were not modeled or excluded as a separate gait-state category in the main ERP analysis. Locomotor state was monitored using the embedded inertial sensors in the EEG system to confirm the sitting and walking conditions, but inertial-sensor-defined epochs such as straight walking, turning, or transient pauses were not analyzed separately. Instead, trials with excessive EEG artifacts were rejected using the same objective preprocessing criteria across conditions. Movement-related contamination was addressed through multiple preprocessing steps, including ICA-based removal of ocular artifacts, sample-level artifact marking, bad-trial rejection, and bad-channel rejection.

#### Permutation test

Permutation tests for time-point and channel-wise comparisons of ERP data between conditions were performed using a two-tailed cluster-based approach implemented with the *ft_timelockstatistics* function in the *FieldTrip* toolbox for MATLAB. Because the sitting and walking conditions were obtained from the same 47 participants, a dependent-samples T-statistic was used at each time point and electrode. Participant identity was specified as the unit variable, and condition was specified as the independent variable in the *FieldTrip* design matrix. For each comparison, condition-specific ERP data were contrasted between the sitting and walking conditions, for example Reg_4-4.06_ in sitting versus Reg_4-4.06_ in walking. Samples exceeding the cluster-defining threshold of *p* < 0.05 were grouped into clusters based on temporal adjacency and spatial proximity among neighbouring electrodes. For the permutation procedure, condition labels were randomly exchanged within each participant, thereby preserving the paired structure of the data. For each of 5000 permutations, a new dependent-samples *t*-value matrix was computed, clusters were formed using the same cluster-defining threshold, and the maximum cluster-level statistic was entered into the null distribution. Observed clusters were considered significant if their cluster-corrected *p*-value was less than 0.05. Tests were performed over the time window from −500 to 2500 ms relative to the transitional click train onset.

### Quantification of EEG response

To quantify onset/change/offset responses, the Response Magnitude (RM) was calculated. The RM is defined as the Root Mean Square (RMS) of the normalized ERP of each electrode within the predefined time window: onset, [0, 300] ms post-onset; change, [50, 350] ms post-change; and offset: [0, 300] ms post-offset). The 300-ms window was selected for both physiological and data-driven reasons. First, the onset, change, and offset responses analyzed here are long-latency auditory cortical ERP responses, whose dominant components occur within the first few hundred milliseconds after an auditory event. A window-based RMS measure provides a robust estimate of response magnitude while reducing sensitivity to peak-latency variability across participants. Second, using the same 300-ms duration for onset, change, and offset responses avoided component-specific window-length bias. Third, the selected windows were supported by the present data because the dominant response energy in the grand-average waveforms and the significant time ranges identified by cluster-based permutation analysis were largely contained within these windows (Song, Xu et al. 2024, Song, Xu et al. 2025, Xu, Huang et al. 2025).

The RM calculation followed the formula:

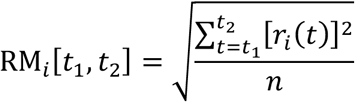

where *r_i_*(*t*) represents the ERP of the *i*^th^ channel at time *t*, *n* represents the number of samples within the time window from *t*_1_ and *t*_2_, and RM*_i_* is the response magnitude of channel *i*. RM was computed separately for each electrode, participant, stimulus condition, motor state (sitting or walking), and task condition (task-engaged or non-task listening). The differential RM (ΔRM), defined as the difference in the response magnitude (RM), was calculated between the two experimental conditions, sitting and walking. Mathematically, it is expressed as:

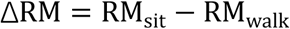

where RM_sit_ is the response magnitude for the sitting condition, and RM_walk_ is the response magnitude for the walking condition. For statistical comparisons between motor states, ΔRM values were averaged across electrodes unless otherwise specified (Figs. 2c–d and 3b–c).

Across all 59 EEG electrodes, TP8 was selected as a representative electrode in a data-driven manner. Specifically, TP8 exhibited the largest change-response ΔRM across all channels for each of the four transitional click trains (Reg_4–4.01_, Reg_4–4.02_, Reg_4–4.03_, and Reg_4–4.06_) in both task-engaged and non-task conditions. For transparency, the location of TP8 is indicated in the topographic maps of ΔRM (Fig. 2c), and example waveforms from this electrode are shown in Fig. 2a. TP8 only served as an example electrode for visualization; statistical inference was performed at the whole-scalp level with multiple-comparisons correction.

### Psychological threshold

The ratio of change detection for each experimental condition was determined by dividing the number of trials in which the participant pressed the right arrow key (indicating perceived change) by the total number of trials in that condition. We fitted psychometric functions for each subject only in Experiment 1 (*n* = 47; Fig. 1c). Psychometric functions were fitted to the data using a cumulative Gaussian model (Yu, Dickman et al. 2015, Xu, Zhai et al. 2017, Du, Song et al. 2024):

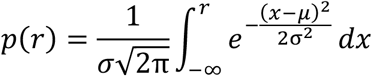

where *p*(*r*) represents the ratio of change detection as a function of ICI *r*. *μ* is the Gaussian mean, and *σ* is the standard deviation (SD). The psychological threshold of change detection was defined as 0.6 of the Gaussian fit (Fig. 1d). This curve fitting procedure was achieved using the ‘psignifit’ software package (see http://bootstrap-software.org/psignifit/) for MATLAB.

### Statistics

To identify channels with significant change responses, the relative Response Magnitude (RM) of the change responses was computed for each channel and subject, as depicted in the topographic plots of RM (Fig. 2b and Fig. 3b). A two-tailed one-sample t-test was performed for each channel and subject, with p-values corrected within channels using the Benjamini-Yekutieli false discovery rate (FDR) procedure (Benjamini and Yekutieli 2001). Channels with significant change responses were marked with black dots on the topographic plots.

For nonparametric tests, the Wilcoxon signed rank test (paired test) and the Mann-Whitney *U* test (unpaired test) was used. The effect size was computed using the r-value.

The r-value was measured and calculated by using the formula:

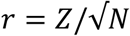

Furthermore, to compute the effect size for each channel, Cohen’s d was calculated using the formula:

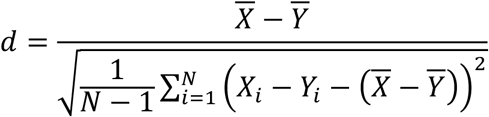

where *d* represents Cohen’s d. *X* and *Y* are the paired samples with a total number of *N* (all *Y* as zero for a one-sample t-test). *X_i_* and *Y_i_* are the *i*^th^ elements from the paired samples. *X̅* and *Y̅* are mean values of the samples.

This method was employed to measure the change response magnitude for each subject and channel in both sitting and walking conditions. The RM was calculated for each subject and condition (sitting and walking) within the predefined time windows for the onset, change, and offset responses

To identify channels with a significant difference in response between correct and incorrect trials in the change detection behavioural task across two experimental conditions, a two-tailed Wilcoxon signed rank test was also performed on the average response relative to the change point, centered around the estimated peak time for each channel.

To test whether there was a clear tuning of RM in the two conditions, a one-way ANOVA was performed among the five contrasted groups (Reg_4-4_, Reg_4-4.01_, Reg_4-4.02_, Reg_4-4.03_, and Reg_4-4.06_). The effect size of the one-way ANOVA was measured using Eta-squared (*η*^2^):

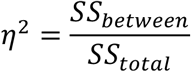

where *SS_between_* is the sum of squared variation due to the factor (between groups). *SS_total_*_<=_ is the total sum of squared variation in the data.

Bayesian statistical analyses were conducted to quantify evidence for the null hypothesis across different types of comparisons. For pairwise comparisons, we used Bayesian paired-sample t-tests with a Jeffreys-Zellner-Siow (JZS) prior, as described by Rouder (Rouder, Speckman et al. 2009). . For group comparisons involving more than two levels, we employed Bayesian one-way ANOVA, using default Cauchy priors on effect sizes (Rouder, Morey et al. 2012). To assess associations between continuous variables, we conducted Bayesian correlation analyses based on a Jeffreys prior for Pearson’s correlation coefficient (Ly, Verhagen et al. 2016). All Bayes factors (BF_10_) were computed using the *BayesFactor* toolbox (Krekelberg 2024) for MATLAB. A BF_10_ greater than 1 indicates evidence in favour of the alternative hypothesis, whereas a BF_10_ less than 1 indicates evidence in favour of the null.

## Data Availability

All data used in this study are available on request.

All codes applied in this study are available on a public Github repository (url: https://github.com:TOMORI233/EEGProcess.git).

## Author contributions

Conceptualization, X.Y.; Methodology, X.Y.; Investigation and Data Curation, I.M., H.X., P.S., and Y.Z.; Writing – Original Draft, X.Y., I.M. and H.X.; Writing – Review & Editing, I.M., H.X., P.S., C.C., Z.D., Y.W., S.G., L.Z., P.C., X.Y., W.W., X.Z., G.C., N.S.P., D.A, M.J.D and X.Y.; Funding Acquisition, X.Y., X.Z. and Y.Z.; Resources, X.Y.; Software, I.M., H.X., P.S., Y.Z., X.B. and H.Y.; Validation and Visualization, I.M. and H.X.; Supervision, X.Y.

## Acknowledgements

We are grateful to Profs. Xiaoming Zhou, Nai Ding and Xi Chen for their invaluable comments on the early version of the manuscript, as well as to Xiaokai Kou for the help with the experiments. This work was supported by Brain Science and Brain-like Intelligence Technology––National Science and Technology Major Project (2022ZD0204800 and 2022ZD0204600) (to X.Y.); National Natural Science Foundation of China 32571216 and 32171044 (to X.Y.), and 32100827 (to Y.Z.); Key Support Discipline Construction Project of Shanghai Municipal Health Commission 2023ZDFC0203 (to X.Z.).

